# Truncated CD19 as a selection marker for the isolation of stem cell derived β-cells

**DOI:** 10.1101/2023.04.05.535733

**Authors:** Luo Ting (Helen) Huang, Dahai Zhang, Cuilan Nian, Lynn Francis C.

## Abstract

Stem cell-derived β-cells (SCβ-cell) are a renewable and scalable alternative to cadaveric islets as a cell replacement therapy for type 1 diabetes (T1D). However, heterogeneity within SCβ-cell cultures remains problematic for graft safety and function. Magnetic selection of SCβ-cells expressing a unique cell surface marker may help deplete undesirable cell types and facilitate functional maturation. Here, we explored CD19 as a potential cell surface marker for the enrichment of insulin-expressing SCβ-cells. Using CRISPR/Cas9 technology, we created a knock-in add-on of CD19-mScarlet downstream of the insulin coding sequence in human embryonic stem cells (hESCs). We established reproducible SCβ-cell surface expression of CD19-mScarlet. Importantly, we developed and optimized a magnetic sorting protocol for CD19-mScarlet-expressing cells, forming enriched SCβ-cell clusters with improved glucose-stimulated c-peptide secretion. This strategy holds promise to facilitate large-scale production of functional SCβ-cells for disease modeling and cell replacement therapy.

## Introduction

Pancreatic β-cells maintain metabolic homeostasis by producing, storing, and releasing insulin in response to fluctuations in blood glucose concentrations. Destruction or dysfunction of pancreatic β-cells leads to insufficient insulin production, chronic hyperglycaemia, and the development of diabetes. While traditional interventions such as exogenous insulin administration can restore normoglycaemia, cadaveric islet transplantation remains one of the most effective treatment options for type 1 diabetes (T1D) (Shapiro, 2012; Shapiro et al., 2017). However, this procedure is limited by the scarce supply and inconsistent quality of islet donors, which poses a barrier to broader applications in a clinical setting (Bellin et al., 2012; Sneddon et al., 2018). Recently, human pluripotent stem cell-derived β-cells (SCβ-cells) have emerged as a renewable, scalable and thus promising alternative for cadaveric islets (Shapiro and Verhoeff, 2023). Over the past decades, several multi-stage differentiation protocols have recapitulated the embryogenesis of pancreatic β-cells from definitive endoderm to pancreatic progenitors, to endocrine progenitors, and pancreatic islets (D’Amour et al., 2006; Hogrebe et al., 2020; Korytnikov and Nostro, 2016; Kroon et al., 2008; Pagliuca et al., 2014; Rezania et al., 2014). Recent efforts have focused on generating mature, INS+ pancreatic β-cells that secrete insulin in response to glucose stimulation (Balboa et al., 2022; Nair et al., 2019; Russ et al., 2015; Velazco-Cruz et al., 2019a). Heterogeneity within SCβ-cell cultures remains problematic, as undesired populations of polyhormonal cells and proliferating pancreatic progenitors may interfere with graft safety and function (Hiyoshi et al., 2022; Robert et al., 2018; Veres et al., 2019).

Enzymatic dissociation and controlled reaggregation have been used to enrich for pancreatic endocrine cells and facilitate maturation (Agulnick et al., 2015; Kelly et al., 2011; Velazco-Cruz et al., 2019). Notably, Nair et al. generated glucose-responsive, enhanced β-clusters by sorting for insulin-expressing cells at the immature β-cell stage (2019). Their method relies on fluorescence-activated cell sorting (FACS) of a transgenic hPSC line, in which an endogenous insulin allele has been replaced by a Green Fluorescent Protein (GFP) reporter (Micallef et al., 2012). FACS limits the scalability of SCβ-cells production, as it entails a time-consuming procedure on a machine that requires extensive calibration and technical expertise; cells are exposed to additional stress and contamination sources during standby.

Alternatively, magnetic sorting of cell surface markers shows promise to deplete undesirable cell types and facilitate differentiation towards mature SCβ-cells. The method has been used to remove human pluripotent stem cells (Fong et al., 2009), isolate anterior definitive endoderm (Mahaddalkar et al., 2020), and enrich for pancreatic progenitors in early stages of differentiation (Ameri et al., 2017; Cogger et al., 2017; Kelly et al., 2011). Sorting for surface markers in early stages of pancreatic differentiation generates heterogenous clusters with relatively higher proportions of INS+ SCβ-cells. On the other hand, sorting for late-stage endocrine cell marker CD49a generated clusters enriched for endocrine cells, but its variable inclusion of α-δ-and pancreatic polypeptide cells (PP-cells) may impede downstream applications such as transplantation (Veres et al., 2019). Recently, Parent et al. identified monoclonal antibodies against immature β-like cells, generating endocrine clusters with improved glucose response and reduced proliferation upon transplantation (Parent et al., 2022). Taken together, magnetic sorting of cell surface markers could offer similar benefits as INS-GFP-based FACS but with greater cost-efficiency and scalability. To capitalize on the best of both worlds, we aimed to generate an hESC line that expresses a cell surface marker for which there are available clinical grade isolation reagents to facilitate a specific and scalable enrichment of insulin-expressing SCβ-cells (Arcangeli et al., 2020)

In this study we explored CD19 as a potential cell surface marker for the enrichment of SCβ-cells. The CD19 antigen is a 95kd transmembrane glycoprotein in the immunoglobulin superfamily. It is expressed across all phases of B cell development, thus commonly used as a biomarker for lymphoma diagnosis and leukemia immunotherapies (Wang et al., 2012). Coincidentally, CD19 is the most frequently targeted antigen in the clinical production of chimeric antigen receptor (CAR) T cells (Arcangeli et al., 2020; Zhang et al., 2018; Zhu et al., 2018). In humans, CD19 is an adaptor protein that recruits cytoplasmic signaling proteins to the cell surface, working with CD21 as a complex to establish the intrinsic threshold for B cell receptor signaling pathways (Tedder et al., 1997). To render a signaling deficient cell surface marker, we truncated the cytoplasmic tail at the C-terminus of CD19 and replaced the intracellular domain with a bright red fluorescent protein mScarlet (Bindels et al., 2016; Wang et al., 2012). Using CRISPR/Cas9 technology, we knocked-in added-on the truncated CD19 downstream of the final insulin coding exon in hESCs. We show that magnetic sorting and reaggregation of immature endocrine cells depleted non-endocrine cells and produced reaggregated SC-β cell clusters that contain up to 70% INS-CD19 expressing cells. Gene expression analyses confirmed that magnetic sorting enriched for INS-expressing cells over other endocrine cell types. Sorted SCβ-cells also showed a trend of increased c-peptide secretion in response to glucose stimulation. By combining CRISPR/Cas9 technology and a scalable magnetic sorting approach, we can facilitate the large-scale production of SCβ-cells for disease modeling and cell replacement therapy.

## Methods

### DNA targeting constructs

A gRNA sequence targeting the 3’ end of the insulin coding sequence (GTGCAACTAGACGCAGCCCGC) was into the pCCC CRISPR/Cas9 vector as previously described to create pCCC-766/767 (Krentz et al., 2014; Novakovsky et al., 2023). For the donor vector we cloned 800bp homology arms into the KpnI/NheI and AscI/NotI sites of Addgene vector #31938 (Hockemeyer et al., 2011). We removed the stop codon, gRNA binding site and PAM sequence from the homology arms to prevent re-cutting of targeted DNA. We next synthesized a Geneblock (IDT DNA Technologies) containing a human codon-optimized version of a PCSK1/3-P2A-CD19Δ-mScarlet and inserted it into the NheI/EcoRI sites to complete the donor vector (Figure S1). The P2A site was included to allow ribosomal skipping and production of membrane-targeted CD19Δ-mScarlet downstream of insulin while the PCSK1/3 site was included to insure removal of any residual 2A protein fragments from the C terminus of insulin within the secretory granule (Kim et al., 2011). CD19Δ-mScarlet contained a signal peptide at the N-terminus for membrane targeting and a series of stop codons at the C terminus but did not include an exogenous polyadenylation sequence. We fully sequenced the cloned regions using Sanger sequencing prior to transfection.

In order to test expression of the codon-optimized version of CD19Δ-mScarlet in HEK293 cells, we subcloned fragments containing only CD19Δ-mScarlet or 2A-CD19Δ-mScarlet into the NheI/EcoRI sites of pCDNA3.1+ (Invitrogen) using PCR.

### HEK293 transfection

To generate transgenic HEK293, cells were seeded at a density of 0.8 x10^6^ cells per well in a six-well suspension plate. 4 µg of the P2A-CD19Δ-mScarlet or CD19Δ-mScarlet construct was transfected using Lipofectamine 2000 (Life Technologies) as per manufacturer’s recommendations. Cells were cultured in DMEM (Gibco) supplemented with 10% fetal bovine serum (Gibco), and Pen/strep (HyClone) in a humidified 5% CO_2_ 37°C incubator for 72 hours before dissociation and staining for flow cytometry analysis.

### Maintenance of human embryonic stem cells

Undifferentiated H1 (WA01; XY) human embryonic stem cells (hESCs) were maintained on Geltrex coated plates (1:100 in DMEM/F12; Gibco) in StemFlex Medium (Gibco) or mTeSR Plus (StemCell Technologies Inc.) in a humidified 5% CO_2,_ 37°C incubator. Cells were passaged every 4 days by non-enzymatic dissociation with ReLeSR (StemCell Technologies Inc.) and plated at a density of 5.0×10^5^ per 60 mm plate.

### Generation of the INS-2A-CD19**Δ**-mScarlet human embryonic stem cell line

To generate the INS-2A-CD19Δ-mScarlet knock-in add-on hESC line, the CRISPR/Cas9 system was used as previously described (Krentz et al., 2014). 4-6 hours prior to electroporation, H1 hESC culture media was replaced with fresh StemFlex (Gibco). hESCs were washed with Ca/Mg-free PBS (PBS-) before dissociation with Accutase for 5 minutes at 37°C. Following detachment, cells were resuspended in Ca/Mg-containing PBS (PBS++) and centrifuged at 200g for 5 minutes. 5×10^6^ cells were resuspended in 0.7ml PBS++, transferred to a 0.4 cm electroporation cuvette with 40 µg of INS-2A-CD19Δ-mScarlet donor plasmid and 15 µg pCCC-LL766/767, and electroporated using Bio-Rad Gene Pulser II system (250 V, 500 µF, infinite resistance, time constant <10 ms). After electroporation, cells were resuspended in Stemflex with 1 mM Y-27632 and plated onto a 60mm Geltrex-coated tissue culture dish. Cells were allowed to recover for up to four days with daily Stemflex media changes before selecting with 0.25 µg/mL puromycin (Sigma Aldrich). Colonies were picked into a 96-well Lamin-521 coated plate (13.3µg/ml in PBS++; Biolamina) within 10 days of electroporation by manually scraping and pipetting the colony off the plate and into a well containing 100 µL of Stemflex with 10 µM Y-27632. Once clones were close to confluent, cells were replica plated onto three plates: one to genotype, one to freeze down and one to expand the correctly targeted clones. Genomic DNA was extracted using QuickExtract (Lucigen) and the following primers, spanning the 5’ and 3’ homology arms were used to genotype: 5’F LL1059: CTTTGAGGCACTGCAAAACTGCGT; 5’R LL688: TGACCGCAGATTCAAGTGTT; 3’F LL789: GAAGGATTGGAGCTACGGGG; 3’R LL790: AGCTCATGGTGCCATCTGAC

To enable clonal selection, the INS-2A-CD19Δ-mScarlet donor plasmid also featured a Lox-P flanked PGK-puro cassette. This was later removed by transfection of Cre-expressing plasmids. Briefly, 4-6 hours prior to electroporation, culture media of INS-2A-CD19Δ-mScarlet hESC clones B3, C4 and E3 were replaced with fresh StemFlex. hESCs were washed with PBS-before dissociation with Accutase for 5 minutes at 37°C. Detached cells were resuspended in PBS++ and centrifuged at 200g for 5 minutes. 5×10^6^ cells were resuspended in 0.7ml PBS++, transferred to a 0.4 cm electroporation cuvette with 15ug of pCAGGS-GFPCre plasmid DNA, and electroporated using Bio-Rad Gene Pulser II system (250 V, 500 µF, infinite resistance, time constants <10 ms). Electroporated cells were resuspended in Stemflex with 1 mM Y-27632 and plated onto a 60mm Geltrex-coated tissue culture dish for recovery.

After 36-48 hours, cells were sorted for GFPCre expression using the BD FACSAria™ II Cell Sorter. 4-6 hours prior to FACS, culture media of hESC clones B3, C4 and E3 were replaced with fresh StemFlex. hESCs were washed with PBS-before dissociation with Accutase for 5 minutes at 37°C. Detached cells were resuspended in 600 μl of PBS++ with ROCK inhibitor and filtered through a 40 µm FACS tube. GFP+ cells were sorted into Stemflex with 1 mM Y-27632 and 1× PenStrep, then plated onto a 60mm Geltrex-coated tissue culture dish. PGK-Puro cassette removal was verified by culturing sorted cells with 0.25µg/ml Puromycin and by genotyping.

### Karyotyping and genomic abnormalities analysis

Genomic integrity was assessed with the hPSC Genetic Analysis Kit (Stemcell, 07550) according to the manufacturer’s instructions. Briefly, 5 ng of genomic DNA was mixed with a ROX reference dye and double-quenched probes tagged with 5-FAM. The probes represented eight common karyotypic abnormalities that have been reported to arise in cloned pluripotent cells: chr 1q, chr 8q, chr 10p, chr 12p, chr 17q, chr 18q, chr 20q or chr Xp. Sample-probe mixes were analyzed on a ViiA^TM^7 PCR System (ThermoFisher Scientific). The results were normalized to the copy number of a control region in chr 4p and analyzed using the ΔΔCt method.

### *In vitro* differentiation of hESCs

To initiate differentiation, we washed 80-90% confluent cultures with PBS-and dissociated cultures into a single-cell suspension with 5 min of Accutase (StemCell Technologies Inc.) treatment at 37°C. Single cells were centrifuged at 200g for 5 minutes and resuspended in StemFlex supplemented with 10 µM Y-27632 (StemCell Technologies Inc.). We seeded 5.5×10^6^ cells per 5.5 mL of mTeSR+ in each well of a six-well suspension plate. The plates were incubated at 37 °C and 5% CO_2_ on an orbital shaker (25mm orbit; Celltron, Infors HT) set at 95 rpm overnight. The next morning, spheroids were washed once with PBS++ (Gibco) and resuspended in day (d) 0 differentiation media with a final volume of 5.5mL per well. Spheroids were fed daily by removing spent media and replenishing with 5.5 mL of fresh media. Media composition for all stages of differentiation is summarized in Supplementary Table 1.

### Magnetic sorting for CD49a and CD19

d25 immature β-like clusters were washed twice with PBS-and dissociated into a single-cell suspension with 8 min of Accutase (StemCell Technologies Inc.) treatment at 37°C. The resulting single cell suspension was diluted in PBS-, centrifuged at 200g for 5 minutes, and resuspended in MACS buffer (PBS-, 0.5%FBS, 2mM EDTA, 10 μM Y-27632). As per manufacturer’s instructions for MACS^®^, cells were filtered through a 30 μm mesh-filter (Miltenyi Biotec), centrifuged, and counted, before staining with mouse anti-CD49a, PE (BD Biosciences, cat# 559596; 1:50) for 30 mins in 4°C with constant agitation. After washing with MACS buffer, cells were resuspended in 300 µL of sorting buffer per 10^7^ cells and anti-PE MicroBeads (Miltenyi Biotec) for 20 min at 4°C with constant agitation. Cells were washed twice and resuspended in 1mL of sorting buffer (up to 10⁸ cells) prior to magnetic separation on LS columns and the MACS Separator (Miltenyi Biotec). Columns were rinsed with 0.5mL of MACS buffer prior to loading the cell suspension. After washing the column of cells three times with MACS buffer, we removed the column from the separator and eluted cells using 1 mL of MACS buffer. The positively selected fractions were reaggregated in Aggrewell^TM^400 (StemCell Technologies Inc.) plates at a concentration of 1.5×10^6^ cells/mL in CB media supplemented with 10 µM Y-27632 and 1× Pen/Strep (Gibco). The plates were incubated at 37 °C and 5% CO_2_ for 72 hours. Later, reaggregated cells were collected into a 6-well non-treated tissue culture plate (Greiner Bio-One) and placed on an orbital shaker set at 100 rpm. Magnetic sorting for CD19 using MACS^®^ followed similar procedures as above, with the exception that dissociated β-like cells were directly incubated with anti-CD19 MicroBeads (Miltenyi Biotec) instead of a primary antibody.

Magnetic sorting for CD19 using the EasySep human CD19 positive selection kit II (StemCell Technologies Inc.) involved the same dissociation procedure as above. Cells were resuspended 10^8^ cells/mL in a 5 mL Falcon round-bottom polystyrene tube (Fisher Scientific). Cells were mixed and incubated with 100 μL/mL of CD19 Positive Selection Cocktail II and 100 μL/mL of Dextran RapidSpheres™ for 3 min respectively. We topped up the tube to 2.5mL and placed it in the EasySep magnet for 3 min before pouring off negatively selected cells by inverting the magnet and tube. The wash procedure was repeated for a total of 3 times.

### Flow cytometry

Day (d)25 immature β-like clusters were washed twice with PBS-and dissociated into a single-cell suspension with 8 min of Accutase (StemCell Technologies Inc.) treatment at 37°C. The single cell suspension was diluted in PBS-, centrifuged at 200g for 5 minutes, resuspended and filtered through a 40 μm mesh-filter FACS tube. Cells were stained with mouse anti-human CD19 antibody (Miltenyi Biotec, cat# 130-113-730; 1:50) in the dark for 25 minutes. Cells were analyzed on a BD LSR II or Fortessa (BCCHRI Flow Core Facility) and data was processed using the FlowJo software.

### Live cell imaging

Unsorted, d25 immature β-like clusters were collected and washed twice with PBS-.

Clusters were stained with 75nM of LysoTracker Blue DND-22 (Invitrogen) in CB media for 30 min. Clusters were co-stained with CellMask Green (Invitrogen; 1:1000) for 5 min before imaging on glass bottom culture plates (MatTek, P35G-1.5-14-C). Images were taken with a 20× oil objective on the Leica TCS SP8 confocal system.

### Intracellular staining

Day 25 immature β-like clusters were washed twice with PBS-and dissociated into a single-cell suspension with 8 min of Accutase (StemCell Technologies Inc.) treatment at 37°C. The single cell suspension was diluted in PBS-, centrifuged at 200g for 5 minutes, resuspended and filtered through a 40 μm mesh-filter FACS tube. To stain for surface CD19 expression, cells were incubated with fixable viability dye eF780 (1:1000) and mouse anti-human CD19 antibody (1:50) for 25 minutes. Cells were fixed overnight at 4°C with the IC Fixation Buffer (eBioscience™). The next day, cells were washed twice with 1× Permeabilization Buffer (eBioscience™) before incubating for 30 min in the same buffer with fixable viability dye eF780 (1:1000) and mouse anti-human CD19 antibody (1:50). Stained cells were washed twice in PBS-before flow cytometry analysis.

### Static glucose stimulated c-peptide secretion assay

50 β-like clusters were loaded into each well of a 24-well plate (Corning) and incubated in low-glucose (2.8mM) KRBH for 1 h in 37°C. Clusters were then transferred to low-glucose and high-glucose (16.7 mM) KRBH for 1 h respectively, and the supernatant was collected. Clusters were then incubated in KRBH containing 16.7 mM glucose and 30 mM KCl (depolarization challenge) for 1 h before the supernatant was collected. Clusters were dispersed using HCl and 70% ethanol to collect total insulin content. Supernatant samples were processed using the STELLUX® Chemi Human C-peptide ELISA (ALPCO).

### Nanostring nCounter XT Gene Expression Assay

50 clusters were lysed in 100μl Buffer RLT (Qiagen) containing 1% β-ME (Sigma Aldrich). Custom nCounter XT Gene Expression Reporter Codeset and Capture Probeset were designed (Supplementary Table 2). Hybridization reactions were set up for nCounter assay as per manufactures instructions. Briefly, 3μl of Reporter Codeset, 5μl of hybridization buffer, 2μl of Capture Probeset, and 1.5μl of total cell lysate was incubated at 65°C for 16 hours. The next morning, a sample cartridge was loaded with 30 μl of each sample and run on the nCounter SPRINT Profiler. Data was normalized to reference genes B2M, GAPDH, GUSB, HPRT1, POLR2A, TBP and analyzed using the nSolver 4.0 Analysis Software.

### Statistical analysis

Statistical analyses were performed using Prism 8 (GraphPad Software). All data are presented as mean ± SEM. Statistically significant differences were assessed using unpaired t-tests or one-way ANOVA followed by post-hoc test for multiple comparisons as appropriate. A p value of < 0.05 was deemed significant.

## Results

### Generation of hESCs with INS-2A-CD19Δ-mScarlet knock-in add-on

To develop a β-cell-specific enrichment strategy that was scalable, accessible, and efficient, we set out to generate an H1 hESC line with expression of a β-cell surface marker, CD19Δ-mScarlet. CD19Δ was fused to mScarlet, a bright red fluorescent protein, to enable downstream analyses (Gadella et al., 2023). We created a knock-in add-on of the CD19Δ-mScarlet downstream of the insulin coding region in H1 hESCs (Figure 1A). To prevent CD19Δ-mScarlet from interfering with endogenous SCβ-cell function, the CD19 was truncated at the C-terminus to render it signaling-deficient (Wang et al., 2012). The knock-in design also featured a 2A site to induce ribosomal skipping during translation of INS and CD19Δ-mScarlet, as well as a proconvertase (PC) 1/3 site to ensure that residual 2A amino acids were cleaved from the endogenous insulin C-terminus within the secretory granules (Liu et al., 2017). Together, the PC 1/3 site and the 2A fragment separate endogenous INS protein expression from cell surface CD19Δ-mScarlet expression. To enable clonal selection, the design also featured a Lox-P flanked PGK-puro cassette which was later removed by transfection of a Cre-expressing plasmid.

**Figure 1.**
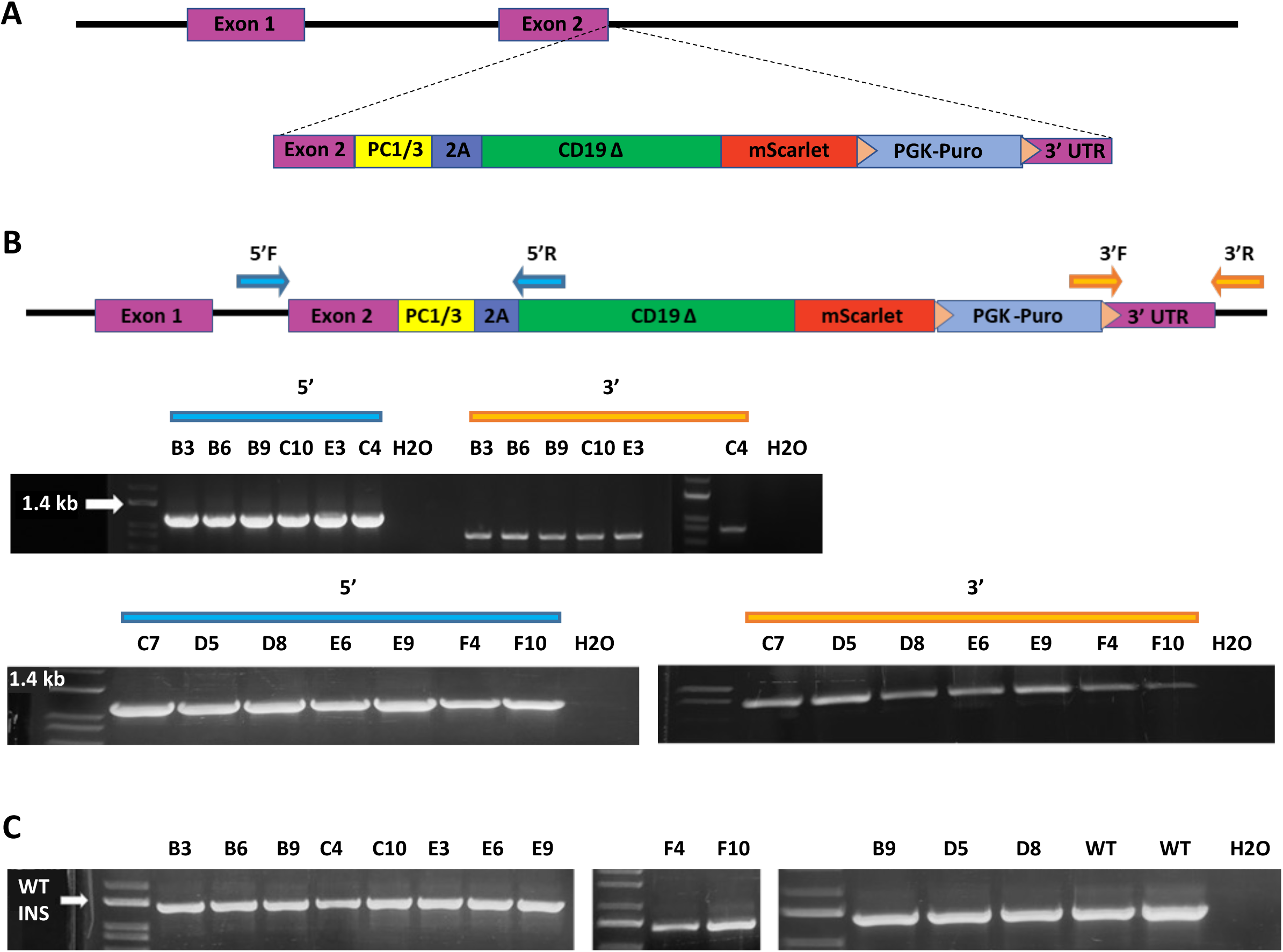
Generation of hESCs with INS-2A-CD19-mScarlet. (A) Schematic overview of the targeting strategy using CRISPR/Cas9 to knock-in add-on 2A-CD19-mScarlet behind the INS coding exon. The DNA donor vector features a proconvertase (PC) 1/3 recognition site, a 2A fragment to facilitate cleavage, and a fusion of signaling-deficient CD19 and mScarlet red fluorescent protein. Regions where genotyping PCR primer pairs bind at 5’ and 3’ highlighted in blue arrows. (B) Genotyping PCR for 3’ and 5’ arms of insertion. (C) Endogenous INS allele assayed with primers 5’F and 3’R for thirteen representative knock-in add-on hESC clones.

Puromycin-resistant clones were genotyped using primer pairs that span the 5’ and 3’ homology arms. 47 out of the 48 clones (98%) were correctly targeted at both the 5’ and 3’ ends as determined using PCR genotyping. 13 clones were expanded, re-genotyped and all of them contained the transgene after expansion (Figure 1B). To characterize whether the insertion targeted one or both alleles, we again used PCR genotyping to distinguish the wild type allele from the modified allele. All 48 clones were heterozygous for the knock-in and all 13 clones identified above remained heterozygous for the CD19Δ-mScarlet knock-in (Figure 1C). Furthermore, sequencing analysis of the PCR products confirmed precise insertion of INS-2A-CD19Δ-mScarlet and that no mutations were introduced near the Cas9 cleavage site (data not shown).

To assess genomic integrity, we performed qPCR-based profiling on genomic hotspot regions that are commonly altered during reprogramming, including 8 common karyotypic abnormalities reported in human ESCs: chromosomes 1q, 4p, 8q, 10p, 12p, 17q, 18q, 20q, Xp. Clone B3, C4 and E3 were karyotypically comparable to the hPSC genomic control, and clone C4 was selected for further differentiation and characterization (Figure S2). In summary, we identified a targeting approach that allows high-efficiency introduction of transgenes downstream of the insulin coding region in human embryonic stem cells. Using this strategy, we generated an INS-PC1/3-2A-CD19Δ-mScarlet hESC line that successfully passed a multi-step quality control workflow.

### CD19-mScarlet localizes to the cell surface of stem cell derived β-cells

In parallel to generating the cell lines we assessed the efficiency of CD19Δ-mScarlet expression and localization to the cell surface. We subcloned the CD19Δ-mScarlet or 2A-CD19Δ-mScarlet sequences into pcDNA3.1 and expressed them using transient transfection on HEK293 cells. On average, 70.5% of transfected HEK293 cells expressed both mScarlet and cell surface CD19Δ, as assessed by flow cytometry (Figure 2A). We found no differences in CD19Δ-mScarlet expression or localization between the two constructs. These experiments demonstrate that the signal peptide in CD19Δ-mScarlet is functional and that it is not affected by the presence of residual 2A sequence at its N terminus.

**Figure 2.**
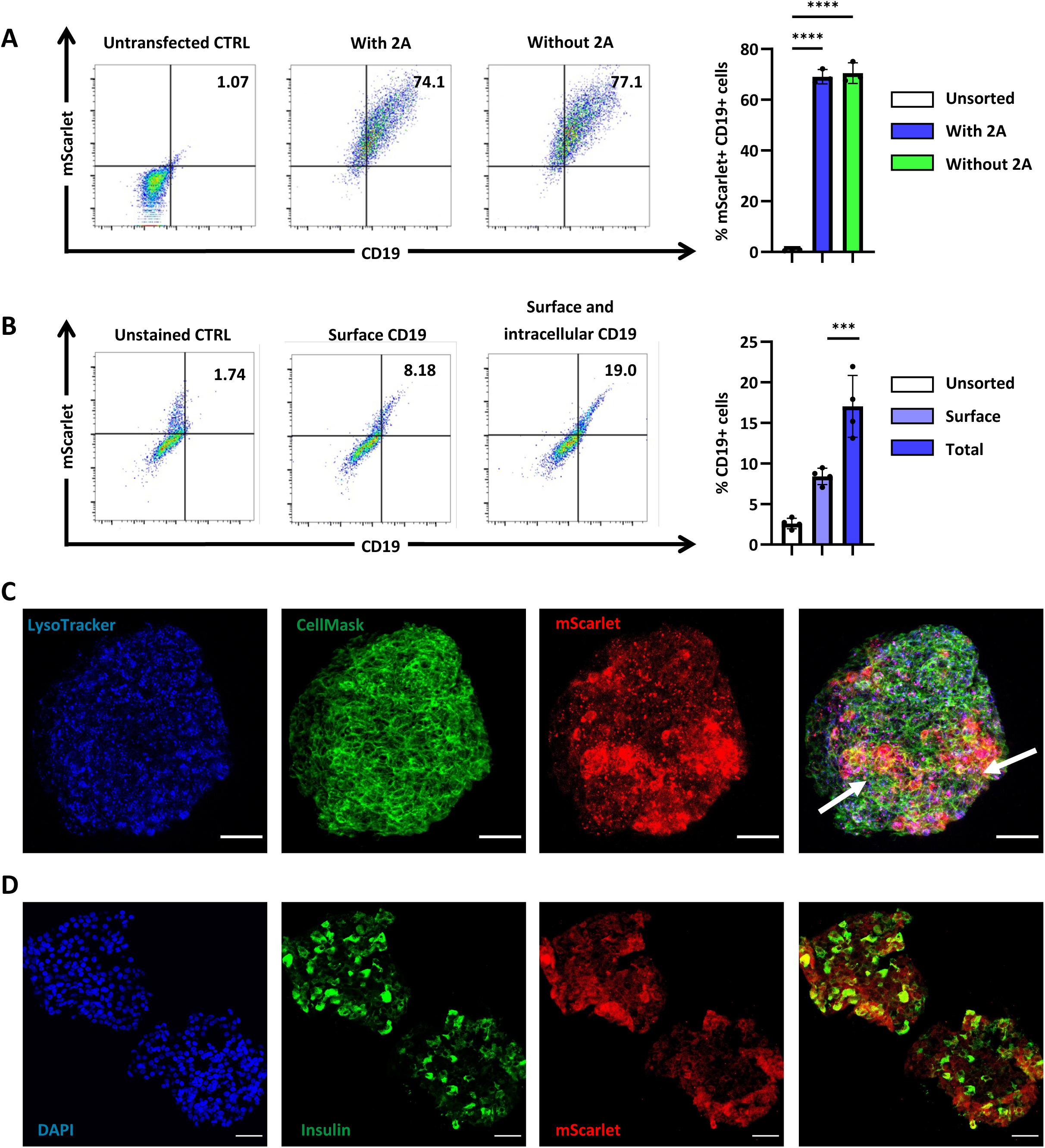
CD19-mScarlet localizes to the cell surface of SCβ-cells. (A) HEK293 cells express CD19Δ-mScarlet 72hrs post transfection. HEK293 cells were transfected with the CD19Δ-mScarlet construct with or without 2A at the N-terminus. Cells were dissociated and assayed for CD19 and mScarlet expression with flow cytometry after 72 hours. N=3 independent biological replicates. (B) Representative flow cytometry analysis of CD19 expression on unstained, surface stained, and surface and intracellularly stained d25 SCβ-cells. N=4 independent biological replicates. ****p<0.0001, ***p<0.001, *p<0.05 by one way ANOVA with multiple comparisons. (C) Live confocal imaging of unsorted stage 6, d25 immature β-like clusters expressing INS-2A-CD19Δ-mScarlet (red). Clusters were stained with LysoTracker (blue) and CellMask (green). Scale bars represent 50 μm. (D) Immunofluorescence analysis of unsorted stage 6, d25 immature β-like clusters expressing INS-2A-CD19Δ-mScarlet (red). Clusters were stained for DAPI (blue) and insulin (green). Scale bars represent 50 μm.

We cultured the C4 hESC clone with INS-2A-CD19Δ-mScarlet knock-in add-on through 25 days and 6 stages of differentiation to derive immature SCβ-cells. To determine whether CD19Δ-mScarlet localizes to the cell surface of SCβ-cells, we compared surface and intracellular CD19 expression in d25 SCβ-cells. Briefly, dissociated SCβ-cells were stained for surface CD19, fixed and permeabilized, then stained again for surface and intracellular CD19. Flow cytometry analysis showed that, in bulk, half of all detectable CD19Δ-mScarlet was expressed on the cell surface (Figure 2B). Live confocal imaging confirmed that CD19Δ-mScarlet co-localizes with the cell membrane in regions of the SCβ-cell cluster that abundantly express CD19Δ-mScarlet (arrows; Figure 2C). Furthermore, immunofluorescence analysis showed insulin expression in CD19Δ-mScarlet-expressing cells of the same cluster (Figure 2D). Interestingly, CD19Δ-mScarlet co-localizes extensively with lysosomes in regions of the cluster that did not abundantly express CD19Δ-mScarlet (Figure 2C). This suggests that SCβ-cells that actively express insulin and CD19Δ-mScarlet traffic a portion of the CD19Δ-mScarlet to the cell surface. In contrast, CD19Δ-mScarlet may be rapidly degraded in stem cell-derived endocrine cells of the cluster where insulin and CD19Δ-mScarlet are not abundantly produced.

### MACS enriches for CD49a on SCβ-cells

We evaluated the efficiency of magnetic sorting by reproducing the enrichment of CD49a-expressing stem cell derived islets cells described by Veres et al. (2019). CD49a sorting entailed an indirect binding approach; CD49a epitopes were bound to CD49a-PE antibodies, which were subsequently bound to anti-PE microbeads for magnetic selection. We used an INS-GFP line that was previously characterized and collected cells for flow cytometry analysis at each stage of magnetic sorting (Novakovsky et al., 2023). Consistent with results reported by Veres et al., magnetically sorted SCβ-cell clusters showed approximately 50% more cells with INS-GFP and surface CD49 expression compared to unsorted stem cell derived islet clusters (Figure 3A&B). An optional enrichment step with a second magnetic column further enriched the INS+ CD49+ proportion by approximately 10% (Figure 3A&B). Recovery rate, as calculated by comparing yield and expected yield of CD49a-expressing cells based on flow cytometry analysis, showed a decreasing trend as the number of column enrichment increases (Figure 3C). The CD49a-sorted cells were seeded in pre-patterned Aggrewells for 3 days and reaggregated into clusters of consistent size (∼100-200um). Cell clusters were transferred to suspension culture for an additional 7 days. After extended culture, the reaggregated SCβ-cells retained high GFP expression (Figure 3D). This suggests that CD49a magnetic sorting enriched for a viable population of INS-GFP-expressing cells, while the presence of microbead attachments did not affect downstream reaggregation of cell clusters.

**Figure 3.**
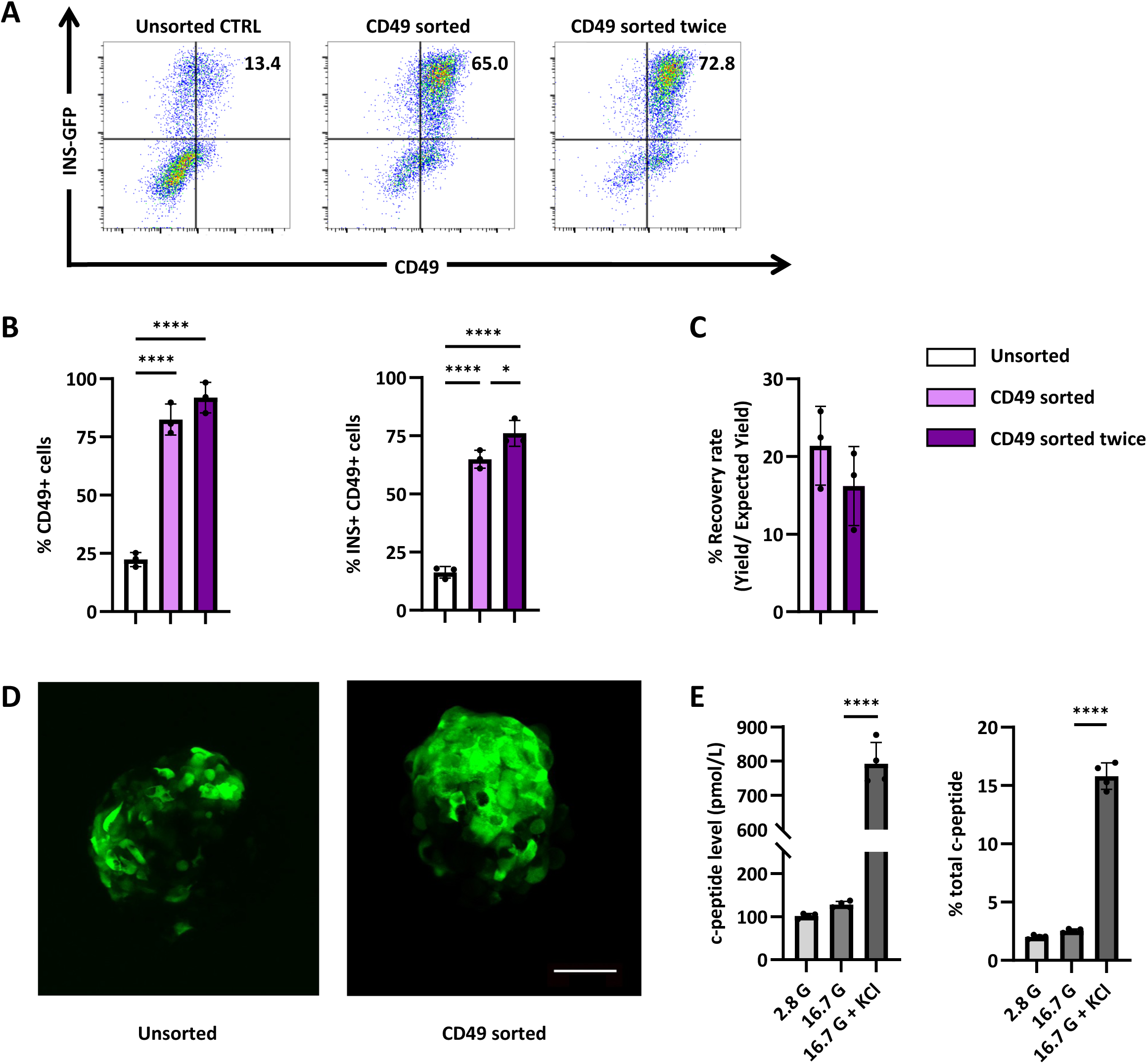
MACS enriches for CD49+ cells in immature INS-GFP SCβ-cells. (A) Representative flow cytometry analysis of CD49 expression on d25 SCβ-cells that were unsorted, CD49 sorted once, and CD49 sorted twice. The proportions of (B) CD49+ cells and CD49+ INS-GFP+ cells were quantified by flow cytometry. N=3 independent biological replicates. ****p<0.0001, *p<0.05 by one way ANOVA with multiple comparisons. (C) Recovery rate was calculated by dividing yield by expected yield. Statistical significance was determined with the unpaired t test. N=3 independent biological replicates. (D) Live confocal imaging of unsorted and sorted stage 6, d35 immature β-like clusters expressing INS-GFP (green). Scale bars represent 50 μm. (E) Static secretion of C-peptide in response to stimulation with 2.8mM glucose, 16.7 mM glucose and 40 mM KCl in an in vitro assay. Total C-peptide was collected by ethanol acid extraction. N=4 independent biological replicates. ****p<0.0001 by one way ANOVA with multiple comparisons.

Reaggregated CD49a-sorted clusters were cultured for 10 days and collected on d35 for functional analysis. To assess the c-peptide secretion response to glucose stimulation, unsorted and CD49a-enriched clusters were incubated in 2.8mM glucose, 16.7mM glucose, and 16.7mM glucose with 30mM KCl for 1 hour consecutively. Supernatants at each stage were collected for c-peptide quantification via ELISA. Compared to unsorted clusters, CD49a-sorted clusters showed limited improvement in glucose-stimulated c-peptide secretion (Figure 3E). Taken together, these results demonstrate that MACS can reproducibly enrich for surface markers expressed on stem cell derived islet cells, which remain viable after extended culture.

### CD19 enriched SCβ-cells have improved INS expression and c-peptide secretory profile

Compared to CD49a sorting, CD19Δ MACS used a direct binding approach, where d25 INS-2A-CD19Δ-mScarlet SCβ-cells were enzymatically dissociated and directly labelled with CD19-specific microbeads. Similar to the pattern previously observed with CD49a sorting, CD19-sorted clusters showed approximately 50% more CD19Δ expression than unsorted SCβ-cell clusters (Figure 4A&B); this trend was consistent in both CD19Δ and mScarlet-expressing populations as expected. The recovery rate of CD19Δ-sorting was comparable to CD49a sorting, both before and after a second optional column enrichment (Figure 4C). We compared commercially available magnetic sorting kits and found that the EasySep CD19 Positive Selection Kit (StemCell Technologies Inc.) offered up to 70% recovery.

**Figure 4.**
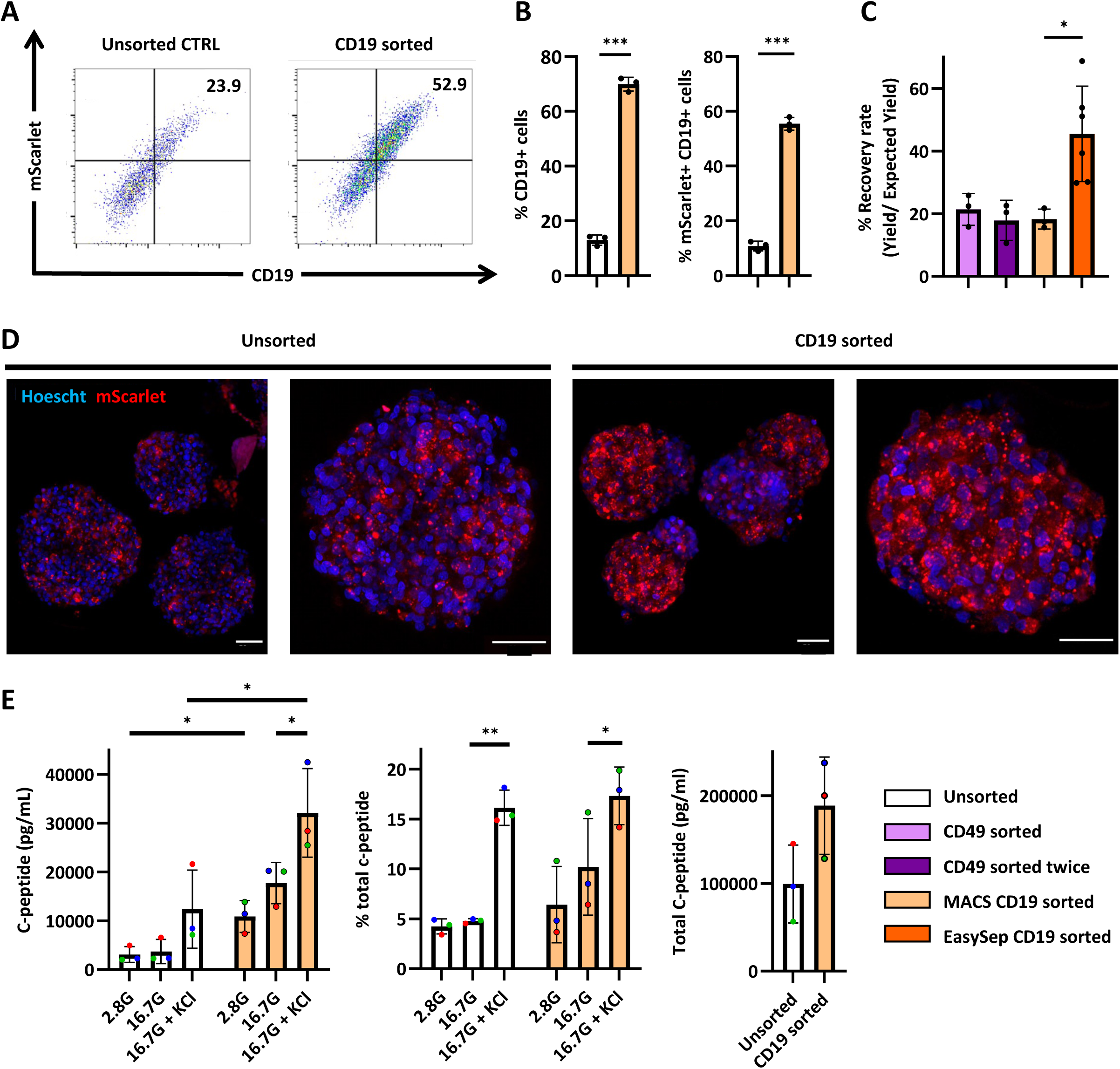
MACS enriches for CD19+ mScarlet+ cells from immature SCβ-cells. (A) Representative flow cytometry analysis of CD19 expression on d35 SCβ-cells that were unsorted and CD19 sorted. The proportions of (B) CD19+ cells and CD19+ mScarlet+ cells were quantified by flow cytometry. N=3 independent biological replicates. ***p<0.001 by one way ANOVA with multiple comparisons. Recovery rate was calculated by dividing yield by expected yield. Statistical significance was determined with the unpaired t test. N=3 independent biological replicates. (C) Live confocal imaging of unsorted and CD19 sorted stage 6, d35 immature β-like clusters expressing INS-2A-CD19-mScarlet (red). Scale bars represent 50 μm. (D) Static secretion of C-peptide in response to stimulation with 2.8mM glucose, 16.7 mM glucose and 40 mM KCl in an in vitro assay. Total C-peptide was collected by ethanol acid extraction. N=3 independent biological replicates. **p<0.01, *p<0.05 by one way ANOVA with multiple comparisons.

The CD19Δ-sorted cells were seeded in pre-patterned Aggrewells for 3 days and reaggregated into clusters of consistent size (∼100-200um). Cell clusters were transferred to suspension culture for an additional 7 days before collecting for live confocal imaging. We visually confirmed that CD19Δ-sorted clusters expressed more mScarlet than unsorted clusters and the mScarlet expression was stable after extended culture (Figure 4D).

To determine whether enrichment for CD19Δ-expressing cells promotes functional maturation, we performed glucose-stimulated C-peptide secretion to test the response of 50 CD19Δ-sorted SCβ-cell clusters to glucose and KCl stimulation. Notably, both basal and total C-peptide secretion increased by approximately 2-fold in CD19Δ-sorted clusters in comparison to unsorted clusters (Figure 4E). This increase correlates with a higher proportion of INS-expressing cells within the clusters, further confirming enrichment of β-cells after sorting. Interestingly, reaggregated CD19Δ-sorted SCβ-cells showed a trend of increased C-peptide secretion in response to 16.7 mM glucose stimulation (Figure 4E).

Gene expression analysis by Nanostring confirmed increased INS expression and decreased GCG and SST expression in CD19Δ-sorted clusters, suggesting that our magnetic sorting strategy enriched for INS-expressing cells over other endocrine cell types (Figure 5). Between CD49a-and CD19Δ-sorting, the latter strategy selected for more INS-expressing cells while depleting more SST, GHRL and PPY-expressing cells. E-cadherin (CDH1), a calcium-dependent cell junction protein that regulates dynamic insulin secretion, was also significantly better enriched by CD19Δ-sorting than CD49a (Dissanayake et al., 2018; Gan et al., 2017). Expression of endocrine markers CHGA and CHGB in sorted clusters both decreased likely due to significant depletion of GCG-expressing cells (Augsornworawat et al., 2020; Veres et al., 2019). Interestingly, CD19Δ-sorted clusters expressed significantly higher levels of IGF2 than unsorted or CD49a-sorted clusters. This may be related to the enrichment of INS-expressing cells, as IGF2 and INS share a promoter and are often expressed as INS-IGF2 hybrid transcripts in the human pancreas (Monk et al., 2006). IGF2 is a potent growth factor recognized for its anti-inflammatory and anti-apoptotic effects within the pancreatic islets, potentially facilitating protective conditions for islet transplantation (Calderari et al., 2007; Hughes et al., 2013; Sandovici et al., 2021). Taken together, these data demonstrate that the scalable magnetic sorting method developed with the INS-2A-CD19Δ-mScarlet hESC line generates enriched SCβ-cell clusters with improved yield, purity, and function over other approaches.

**Figure 5.**
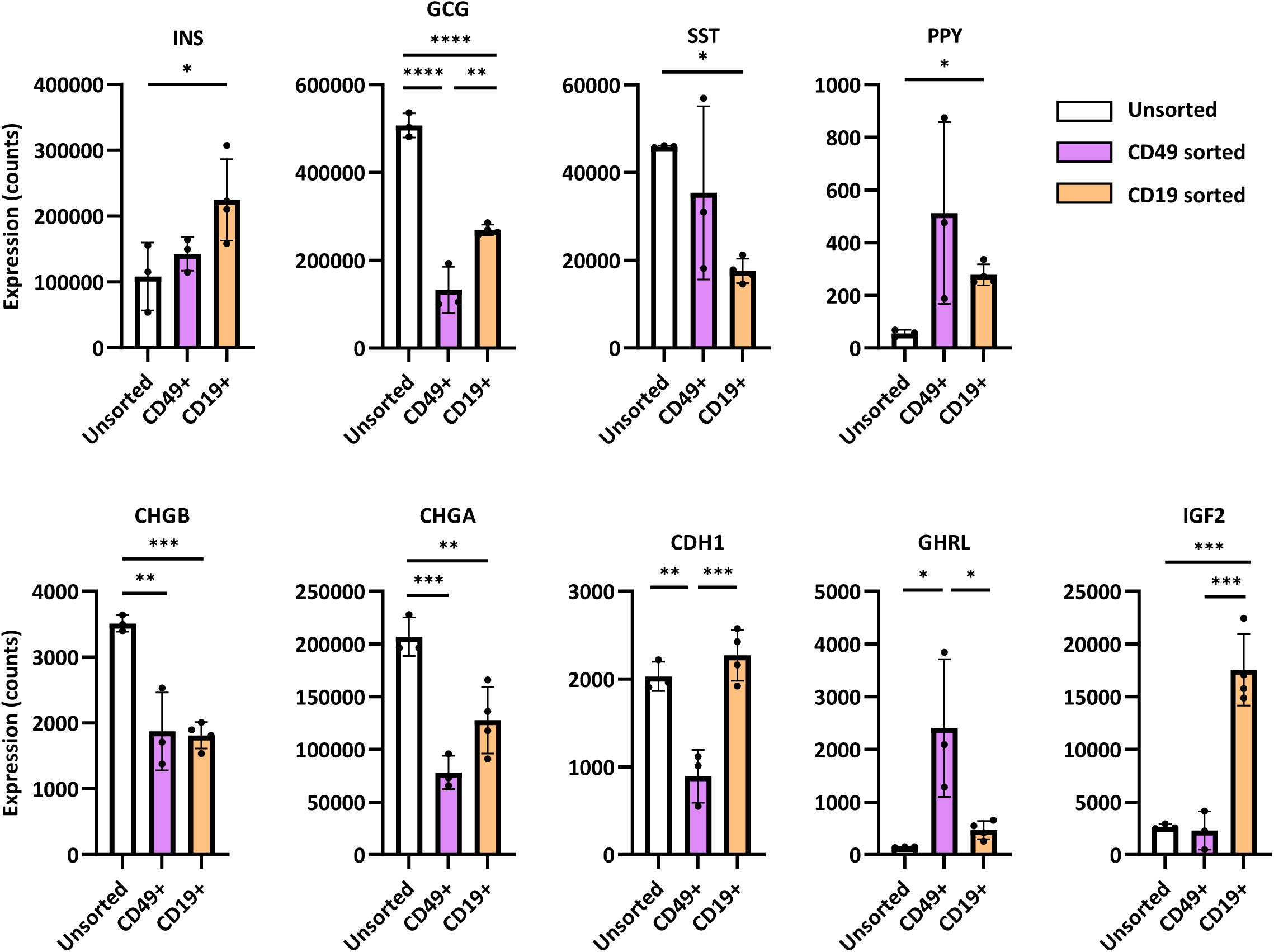
Gene expression analysis of unsorted, CD49 sorted and CD19 sorted SCβ-cells with Nanostring on the following genes: INS, GCG, SST, PPY, CHGB, CHGA, GHRL, CDH1, and IGF2. N=3 independent biological replicates.

## Discussion

SCβ-cell replacement therapy with is a promising alternative to existing treatment options for T1D. However, heterogeneity within SCβ-cell cultures remains problematic for graft safety and long-term function. Here, we developed a scalable and SCβ-cell-specific selection protocol that does not rely on fluorescent reporters of insulin expression. Using CRISPR/Cas9, we knocked-in added-on a truncated form of CD19 downstream of the insulin coding exon in hESCs. We confirmed reproducible cell surface expression of CD19Δ-mScarlet. Importantly, we developed and optimized a magnetic sorting method for INS-CD19Δ-expressing cells, forming enriched SCβ-cell clusters.

To generate a reporter cell line that faithfully reflects INS expression, Nair et al. used an INS^GFP/W^ line that replaced one of the INS alleles with GFP (Micallef et al., 2012). In contrast, our INS-2A-CD19Δ-mScarlet cell line featured a heterozygous knock-in add-on. The DNA vector includes a PC1/3 site and a 2A fragment to separate endogenous INS activity from cell surface CD19Δ-mScarlet expression. This design allows both copies of the INS allele to retain endogenous function, presumably enabling the reaggregated SCβ-cell to mature functionally and respond more physiologically to external stimulus.

Intracellular staining for CD19Δ shows that only half of all detectable CD19Δ in the SCβ-cell is on the cell surface. One possible explanation is that, even with the incorporation of PC1/3 and 2A in the donor construct, a proportion of CD19Δ-mScarlet remain did not fold properly and was targeted to the lysosomal compartment for degradation (Figure 2C). Although we confirmed stable and efficient cell-surface expression of CD19Δ-mScarlet in HEK293 cells, we cannot exclude the possibility that improperly folded CD19Δ-mScarlet may be targeted to lysosomes in immature SCβ-cells. Future iterations of the INS-2A-CD19Δ-mScarlet cell line may incorporate the addition of a few glycine-serine linkers between CD19Δ and mScarlet to allow more of the CD19Δ to properly fold and target the cell membrane (Van Rosmalen et al., 2017).

Previous studies reported magnetic sorting strategies that enrich for anterior definitive endoderm (Mahaddalkar et al., 2020) and pancreatic progenitors (Ameri et al., 2017; Cogger et al., 2017; Kelly et al., 2011). Sorting for surface markers in early stages of pancreatic differentiation generated SCβ-cell clusters that remain heterogenous with less than 60% of insulin-expressing cells. On the other hand, sorting for endocrine cell marker CD49a generated clusters that contain up to 80% SCβ-cells with improved response to glucose *in vitro* (Veres et al., 2019). In our hands, CD49a sorting generated clusters with approximately 65% INS-GFP and CD49a-expressing cells but these cells showed limited c-peptide response to glucose stimulation.

In contrast to endogenous endocrine markers, such as CD49a, we chose to sort for a surface marker that is specific for INS-expressing cells. We observed a trend of improvement in glucose-stimulated-c-peptide secretion from CD19Δ-sorted clusters. Gene expression analyses further indicate that CD19Δ-sorting was more effective than CD49a at enriching for INS-expressing cells and depleting the SST, PPY, and GHRL-expressing populations. Despite significant enrichment, both CD49a and CD19Δ-sorted clusters remained heterogenous, containing approximately 30% INS-cells. This may be related to the inclusion of polyhormonal cells that express *INS* temporarily before resolving into monohormonal GCG or SST-expressing cells upon further maturation (Alvarez-Dominguez et al., 2020). Future attempts at magnetic sorting may consider extending initial culture time to 35 days, when a large majority of polyhormonal cells have resolved into more defined cell types.

In order to further enrich for INS-expressing SCβ-cells, sequential magnetic selections may be employed at the expense of cell yield and recovery (Figure 4C). The sequential separations may feature two rounds of positive CD19Δ enrichment. They may also positively select for two different SCβ-cell markers of interest, such as a combination of CD49a labelling followed by CD19Δ labelling. Alternatively, the separation may start by negatively selecting for non-SCβ-cell surface markers, such as GP2 (Ameri et al., 2017), then following with CD19Δ-based positive selection. This latter approach is preferrable because it does not require additional manipulation to release and deplete magnetic beads between sequential steps.

In summary, we developed a SCβ-cell-specific selection protocol and optimized scalable enrichment conditions to facilitate efficient recovery of INS-expressing cells from hESC pancreatic differentiations. Magnetic sorting of genetically engineered cell surface markers should therefore allow the large-scale production of functional islet-like clusters suitable for disease modeling and cell replacement therapies for patients with T1D.

## Acknowledgements

The authors thank the members of the Lynn, Levings, Verchere, Wasserman and Johnson Laboratories (Vancouver, British Columbia, Canada); the Bruin Laboratory (Carleton University) and the PE MacDonald Laboratory (University of Alberta) for technical support, discussion, and critical reading of the manuscript. F.C.L. was supported by the JDRF (5-SRA-2020-1059-S-B, 3-COE-2022-1103-M-B) and Canadian Institutes of Health Research (ASD-173663). Salary (F.C.L.) was supported by the Michael Smith Foundation for Health Research (#5238 BIOM) and the BC Children’s Hospital Research Institute. Canada Graduate Scholarship support was provided by the National Science and Engineering Research Council (6563; L.T.H). The authors acknowledge that UBC and BC Children’s Hospital are situated on the traditional, ancestral, and unceded territories of the Coast Salish peoples, the Sḵwx̱ wú7mesh (Squamish), səl^̓^ ilwətaɁɬ (Tsleil-Waututh), and xʷməθkʷəy̓ əm (Musqueam) Nations.

---

**Figure S1.**
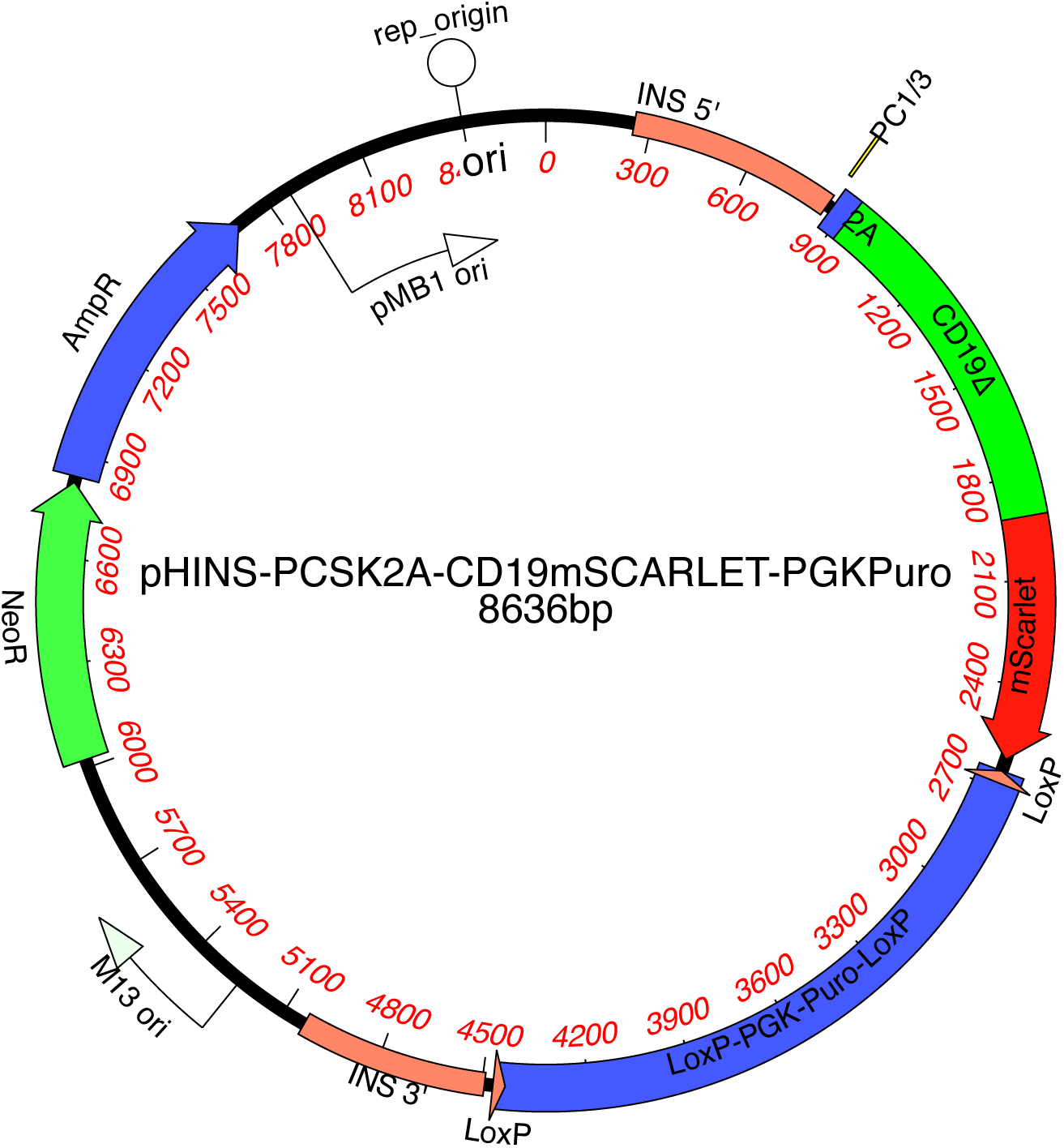
Schematic representation of the complete DNA donor vector, featuring 800bp of INS homology arms and the human codon-optimized version of PCSK1/3-P2A-CD19Δ-mScarlet.

**Figure S2.**
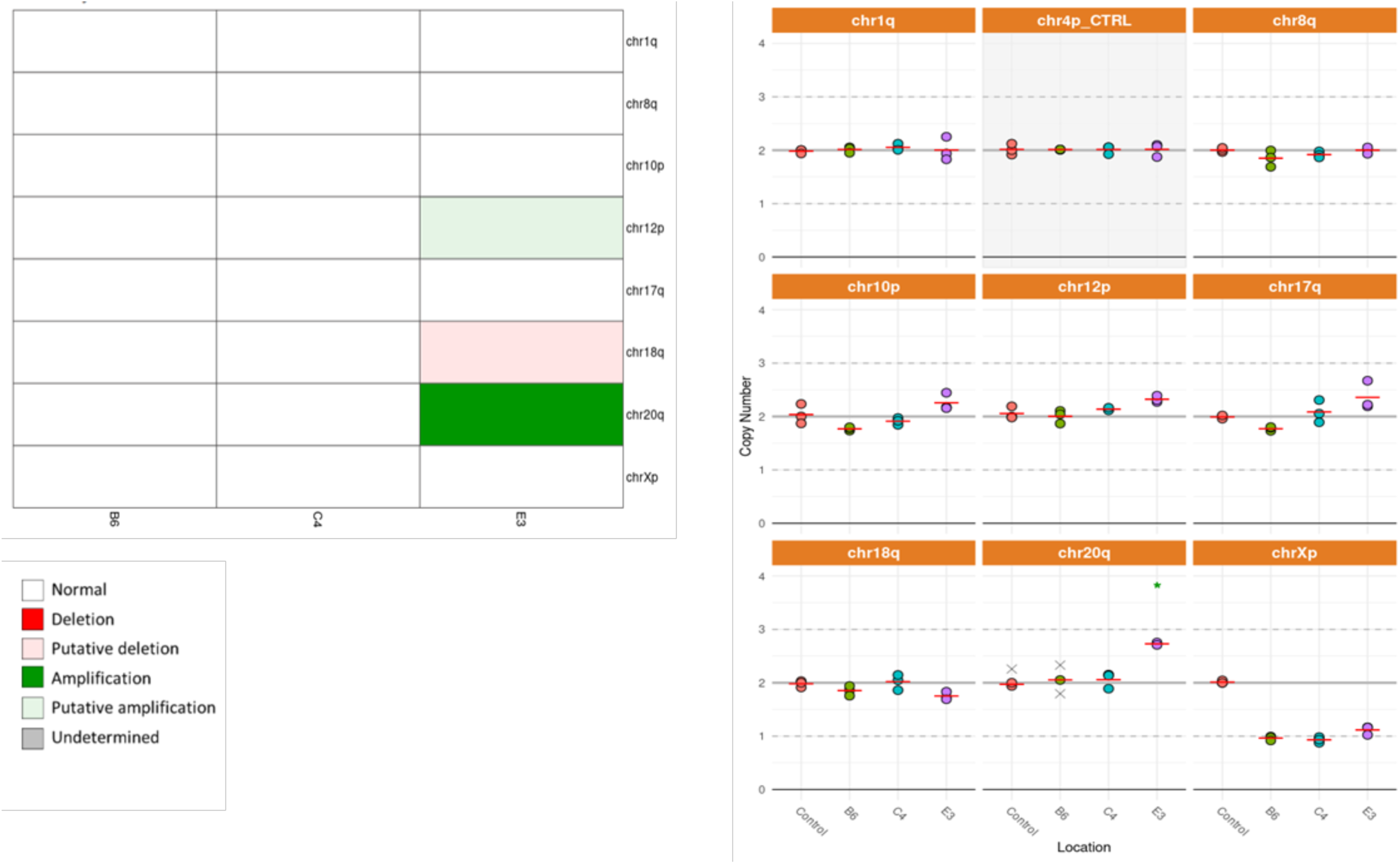
qPCR-based profiling of genomic abnormalities in INS-2A-CD19-mScarlet cell clones B6, C4 and E3. N=3 technical replicates. The probes represent eight common karyotypic abnormalities that have been reported to arise in hESC and hiPSC: chr 1q, chr 8q, chr 10p, chr 12p, chr 17q, chr 18q, chr 20q or chr Xp. Sample-probe mixes were analyzed on a ViAATM7 PCR System. Results were normalized to the copy number of a control region in chr 4p and analyzed using the ΔΔCt method.

**Figure S3.**
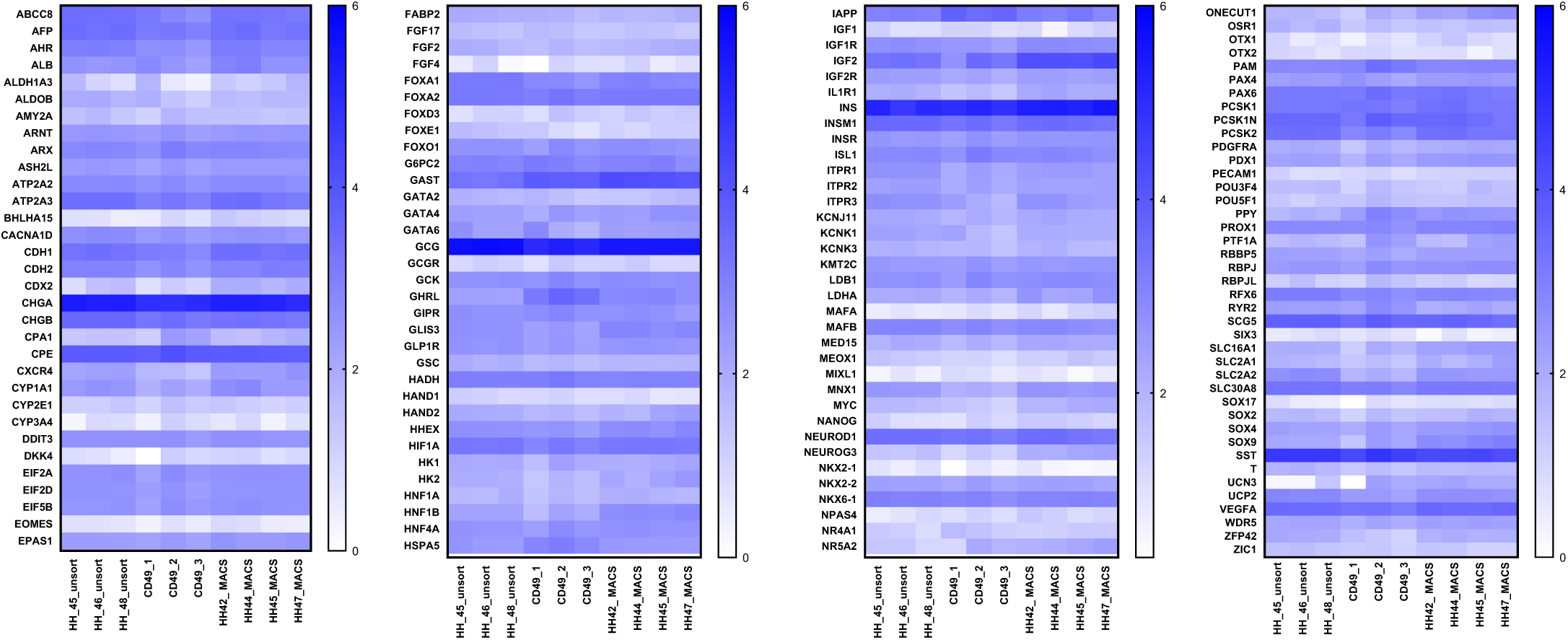
Gene expression analysis of unsorted, CD49 sorted and CD19 sorted SCβ-cells with Nanostring. 50 clusters were lysed in buffer RLT and 1% β-ME. Hybridization reactions were set up for nCounter assay with custom designed XT Gene Expression Reporter Codeset and Capture Probesets. Data was normalized to reference genes and analyzed using the nSolver 4.0 Analysis Software.

## References

Agulnick, et al. (2015). Insulin-Producing Endocrine Cells Differentiated In Vitro From Human Embryonic Stem Cells Function in Macroencapsulation Devices In Vivo. 1214–1222.

Alvarez-Dominguez, J.R., Donaghey, J., Rasouli, N., Kenty, J.H.R., Helman, A., Charlton, J., Straubhaar, J.R., Meissner, A., and Melton, D.A. (2020). Circadian Entrainment Triggers Maturation of Human In Vitro Islets. Cell Stem Cell 26, 108–122.e10. https://doi.org/10.1016/j.stem.2019.11.011.

Ameri, J., Borup, R., Prawiro, C., Ramond, C., Schachter, K.A., Scharfmann, R., and Semb, H. (2017). Efficient Generation of Glucose-Responsive Beta Cells from Isolated GP2+ Human Pancreatic Progenitors. Cell Rep 19, 36–49. https://doi.org/10.1016/j.celrep.2017.03.032.

Arcangeli, S., Falcone, L., Camisa, B., De Girardi, F., Biondi, M., Giglio, F., Ciceri, F., Bonini, C., Bondanza, A., and Casucci, M. (2020). Next-Generation Manufacturing Protocols Enriching TSCM CAR T Cells Can Overcome Disease-Specific T Cell Defects in Cancer Patients. Front Immunol 11. https://doi.org/10.3389/fimmu.2020.01217.

Augsornworawat, P., Maxwell, K.G., Velazco-Cruz, L., and Millman, J.R. (2020). Single-Cell Transcriptome Profiling Reveals β Cell Maturation in Stem Cell-Derived Islets after Transplantation. Cell Rep 32. https://doi.org/10.1016/j.celrep.2020.108067.

Balboa, D., Barsby, T., Lithovius, V., Saarimäki-Vire, J., Omar-Hmeadi, M., Dyachok, O., Montaser, H., Lund, P.E., Yang, M., Ibrahim, H., et al. (2022). Functional, metabolic and transcriptional maturation of human pancreatic islets derived from stem cells. Nat Biotechnol 40, 1042–1055. https://doi.org/10.1038/s41587-022-01219-z.

Bellin, M.D., Barton, F.B., Heitman, A., Harmon, J. V., Kandaswamy, R., Balamurugan, A.N., Sutherland, D.E.R., Alejandro, R., and Hering, B.J. (2012). Potent induction immunotherapy promotes long-term insulin independence after islet transplantation in type 1 diabetes. American Journal of Transplantation 12, 1576–1583. https://doi.org/10.1111/j.1600-6143.2011.03977.x.

Bindels, D.S., Haarbosch, L., Van Weeren, L., Postma, M., Wiese, K.E., Mastop, M., Aumonier, S., Gotthard, G., Royant, A., Hink, M.A., et al. (2016). MScarlet: A bright monomeric red fluorescent protein for cellular imaging. Nat Methods 14, 53–56. https://doi.org/10.1038/nmeth.4074.

Calderari, S., Gangnerau, M.N., Thibault, M., Meile, M.J., Kassis, N., Alvarez, C., Portha, B., and Serradas, P. (2007). Defective IGF2 and IGF1R protein production in embryonic pancreas precedes beta cell mass anomaly in the Goto-Kakizaki rat model of type 2 diabetes. Diabetologia 50, 1463–1471. https://doi.org/10.1007/s00125-007-0676-2.

Cogger, K.F., Sinha, A., Sarangi, F., McGaugh, E.C., Saunders, D., Dorrell, C., Mejia-Guerrero, S., Aghazadeh, Y., Rourke, J.L., Screaton, R.A., et al. (2017). Glycoprotein 2 is a specific cell surface marker of human pancreatic progenitors. Nat Commun 8. https://doi.org/10.1038/s41467-017-00561-0.

D’Amour, K.A., Bang, A.G., Eliazer, S., Kelly, O.G., Agulnick, A.D., Smart, N.G., Moorman, M.A., Kroon, E., Carpenter, M.K., and Baetge, E.E. (2006). Production of pancreatic hormone-expressing endocrine cells from human embryonic stem cells. Nat Biotechnol 24, 1392–1401. https://doi.org/10.1038/nbt1259.

Dissanayake, W.C., Sorrenson, B., and Shepherd, P.R. (2018). The role of adherens junction proteins in the regulation of insulin secretion. Biosci Rep 38. https://doi.org/10.1042/BSR20170989.

Fong, C.Y., Peh, G.S.L., Gauthaman, K., and Bongso, A. (2009). Separation of SSEA-4 and TRA-1-60 labelled undifferentiated human embryonic stem cells from a heterogeneous cell population using magnetic-activated cell sorting (MACS) and fluorescence-activated cell sorting (FACS). Stem Cell Rev Rep 5, 72–80. https://doi.org/10.1007/s12015-009-9054-4.

Gadella, T.W.J., van Weeren, L., Stouthamer, J., Hink, M.A., Wolters, A.H.G., Giepmans, B.N.G., Aumonier, S., Dupuy, J., and Royant, A. (2023). mScarlet3: a brilliant and fast-maturing red fluorescent protein. Nat Methods https://doi.org/10.1038/s41592-023-01809-y.

Gan, W.J., Zavortink, M., Ludick, C., Templin, R., Webb, R., Webb, R., Ma, W., Poronnik, P., Parton, R.G., Gaisano, H.Y., et al. (2017). Cell polarity defines three distinct domains in pancreatic β-cells. J Cell Sci 130, 143–151. https://doi.org/10.1242/jcs.185116.

Hiyoshi, H., Sakuma, K., Tsubooka-Yamazoe, N., Asano, S., Mochida, T., Yamaura, J., Konagaya, S., Fujii, R., Matsumoto, H., Ito, R., et al. (2022). Characterization and reduction of non-endocrine cells accompanying islet-like endocrine cells differentiated from human iPSC. Sci Rep 12. https://doi.org/10.1038/s41598-022-08753-5.

Hockemeyer, D., Wang, H., Kiani, S., Lai, C.S., Gao, Q., Cassady, J.P., Cost, G.J., Zhang, L., Santiago, Y., Miller, J.C., et al. (2011). Genetic engineering of human pluripotent cells using TALE nucleases. Nat Biotechnol 29, 731–734. https://doi.org/10.1038/nbt.1927.

Hogrebe, N.J., Augsornworawat, P., Maxwell, K.G., Velazco-Cruz, L., and Millman, J.R. (2020). Targeting the cytoskeleton to direct pancreatic differentiation of human pluripotent stem cells. Nat Biotechnol 97–115. https://doi.org/10.1038/s41587-020-0430-6.

Hughes, A., Mohanasundaram, D., Kireta, S., Jessup, C.F., Drogemuller, C.J., and Coates, P.T.H. (2013). Insulin-like growth factor-II (IGF-II) prevents proinflammatory cytokine-induced apoptosis and significantly improves islet survival after transplantation. Transplantation 95, 671–678. https://doi.org/10.1097/TP.0b013e31827fa453.

James Shapiro, A.M. (2012). Islet transplantation in type 1 diabetes: Ongoing challenges, refined procedures, and long-term outcome. Review of Diabetic Studies 9, 385–406. https://doi.org/10.1900/RDS.2012.9.385.

Kelly, O.G., Chan, M.Y., Martinson, L.A., Kadoya, K., Ostertag, T.M., Ross, K.G., Richardson, M., Carpenter, M.K., D’Amour, K.A., Kroon, E., et al. (2011). Cell-surface markers for the isolation of pancreatic cell types derived from human embryonic stem cells. Nat Biotechnol 29, 750–756. https://doi.org/10.1038/nbt.1931.

Kim, J.H., Lee, S.R., Li, L.H., Park, H.J., Park, J.H., Lee, K.Y., Kim, M.K., Shin, B.A., and Choi, S.Y. (2011). High cleavage efficiency of a 2A peptide derived from porcine teschovirus-1 in human cell lines, zebrafish and mice. PLoS One 6. https://doi.org/10.1371/journal.pone.0018556.

Korytnikov, R., and Nostro, M.C. (2016). Generation of polyhormonal and multipotent pancreatic progenitor lineages from human pluripotent stem cells. Methods 101, 56–64. https://doi.org/10.1016/j.ymeth.2015.10.017.

Krentz, N.A.J., Nian, C., and Lynn, F.C. (2014). TALEN/CRISPR-mediated eGFP knock-in add-on at the OCT4 locus does not impact differentiation of human embryonic stem cells towards endoderm. PLoS One 9, 1–21. https://doi.org/10.1371/journal.pone.0114275.

Kroon, E., Martinson, L.A., Kadoya, K., Bang, A.G., Kelly, O.G., Eliazer, S., Young, H., Richardson, M., Smart, N.G., Cunningham, J., et al. (2008). Pancreatic endoderm derived from human embryonic stem cells generates glucose-responsive insulin-secreting cells in vivo. Nat Biotechnol 26, 443–452. https://doi.org/10.1038/nbt1393.

Liu, Z., Chen, O., Wall, J.B.J., Zheng, M., Zhou, Y., Wang, L., Ruth Vaseghi, H., Qian, L., and Liu, J. (2017). Systematic comparison of 2A peptides for cloning multi-genes in a polycistronic vector. Sci Rep 7, 1–9. https://doi.org/10.1038/s41598-017-02460-2.

Mahaddalkar, P.U., Scheibner, K., Pfluger, S., Ansarullah, Sterr, M., Beckenbauer, J., Irmler, M., Beckers, J., Knöbel, S., and Lickert, H. (2020). Generation of pancreatic β cells from CD177+ anterior definitive endoderm. Nat Biotechnol 38, 1061–1072. https://doi.org/10.1038/s41587-020-0492-5.

Micallef, S.J., Li, X., Schiesser, J. v., Hirst, C.E., Yu, Q.C., Lim, S.M., Nostro, M.C., Elliott, D.A., Sarangi, F., Harrison, L.C., et al. (2012). INSGFP/w human embryonic stem cells facilitate isolation of in vitro derived insulin-producing cells. Diabetologia 55, 694–706. https://doi.org/10.1007/s00125-011-2379-y.

Monk, D., Sanches, R., Arnaud, P., Apostolidou, S., Hills, F.A., Abu-Amero, S., Murrell, A., Friess, H., Reik, W., Stanier, P., et al. (2006). Imprinting of IGF2 P0 transcript and novel alternatively spliced INS-IGF2 isoforms show differences between mouse and human. Hum Mol Genet 15, 1259–1269. https://doi.org/10.1093/hmg/ddl041.

Nair, G.G., Liu, J.S., Russ, H.A., Tran, S., Saxton, M.S., Chen, R., Juang, C., Li, M. lan Nguyen, V.Q., Giacometti, S., et al. (2019). Recapitulating endocrine cell clustering in culture promotes maturation of human stem-cell-derived β cells. Nat Cell Biol 21, 263–274. https://doi.org/10.1038/s41556-018-0271-4.

Novakovsky, G., Sasaki, S., Fornes, O., Omur, M.E., Huang, H., Bayly, C.L., Zhang, D., Lim, N., Cherkasov, A., Pavlidis, P., et al. (2023). In silico discovery of small molecules for efficient stem cell differentiation into definitive endoderm. Stem Cell Reports 18, 765– 781. https://doi.org/10.1016/j.stemcr.2023.01.008.

Pagliuca, F.W., Millman, J.R., Gürtler, M., Segel, M., Van Dervort, A., Ryu, J.H., Peterson, Q.P., Greiner, D., and Melton, D.A. (2014). Generation of functional human pancreatic β cells in vitro. Cell 159, 428–439. https://doi.org/10.1016/j.cell.2014.09.040.

Parent, A. v., Ashe, S., Nair, G.G., Li, M.L., Chavez, J., Liu, J.S., Zhong, Y., Streeter, P.R., and Hebrok, M. (2022). Development of a scalable method to isolate subsets of stem cell-derived pancreatic islet cells. Stem Cell Reports 17, 979–992. https://doi.org/10.1016/j.stemcr.2022.02.001.

Rezania, A., Bruin, J.E., Arora, P., Rubin, A., Batushansky, I., Asadi, A., Dwyer, S.O., Quiskamp, N., Mojibian, M., Albrecht, T., et al. (2014). articles Reversal of diabetes with insulin-producing cells derived in vitro from human pluripotent stem cells. 32, 1121–1134. https://doi.org/10.1038/nbt.3033.

Robert, T., de Mesmaeker, I., Stangé, G.M., Suenens, K.G., Ling, Z., Kroon, E.J., and Pipeleers, D.G. (2018). Functional Beta Cell Mass from Device-Encapsulated hESC-Derived Pancreatic Endoderm Achieving Metabolic Control. Stem Cell Reports 10, 739–750. https://doi.org/10.1016/j.stemcr.2018.01.040.

Van Rosmalen, M., Krom, M., and Merkx, M. (2017). Tuning the Flexibility of Glycine-Serine Linkers to Allow Rational Design of Multidomain Proteins. Biochemistry 56, 6565– 6574. https://doi.org/10.1021/acs.biochem.7b00902.

Russ, H.A., Parent, A. V, Ringler, J.J., Hennings, T.G., Nair, G.G., Shveygert, M., Guo, T., Puri, S., Haataja, L., Cirulli, V., et al. (2015). Controlled induction of human pancreatic progenitors produces functional beta-like cells in vitro. EMBO J 34, 1759–1772. https://doi.org/10.15252/embj.201591058.

Sandovici, I., Hammerle, C.M., Virtue, S., Vivas-Garcia, Y., Izquierdo-Lahuerta, A., Ozanne, S.E., Vidal-Puig, A., Medina-Gómez, G., and Constância, M. (2021). Autocrine IGF2 programmes β-cell plasticity under conditions of increased metabolic demand. Sci Rep 11. https://doi.org/10.1038/s41598-021-87292-x.

Shapiro, A.M.J., and Verhoeff, K. (2023). A spectacular year for islet and stem cell transplantation. Nat Rev Endocrinol 19, 68–69. https://doi.org/10.1038/s41574-022-00790-4.

Shapiro, A.M.J., Pokrywczynska, M., and Ricordi, C. (2017). Clinical pancreatic islet transplantation. Nat Rev Endocrinol 13, 268–277. https://doi.org/10.1038/nrendo.2016.178.

Sneddon, J.B., Tang, Q., Stock, P., Bluestone, J.A., Roy, S., Desai, T., and Hebrok, M. (2018). Stem Cell Therapies for Treating Diabetes: Progress and Remaining Challenges. Cell Stem Cell 22, 810–823. https://doi.org/10.1016/j.stem.2018.05.016.

Tedder, T.F., Inaoki, M., and Sato, S. (1997). The CD19-CD21 complex regulates signal transduction thresholds governing humoral immunity and autoimmunity. Immunity 6, 107–118. https://doi.org/10.1016/S1074-7613(00)80418-5.

Velazco-Cruz, L., Song, J., Maxwell, K.G., Goedegebuure, M.M., Augsornworawat, P., Hogrebe, N.J., and Millman, J.R. (2019a). Acquisition of Dynamic Function in Human Stem Cell-Derived β Cells. Stem Cell Reports 12, 351–365. https://doi.org/10.1016/j.stemcr.2018.12.012.

Velazco-Cruz, L., Song, J., Maxwell, K.G., Goedegebuure, M.M., Augsornworawat, P., Hogrebe, N.J., and Millman, J.R. (2019b). Acquisition of Dynamic Function in Human Stem Cell-Derived β Cells. Stem Cell Reports 12, 351–365. https://doi.org/10.1016/j.stemcr.2018.12.012.

Veres, A., Faust, A.L., Bushnell, H.L., Engquist, E.N., Kenty, J.H.R., Harb, G., Poh, Y.C., Sintov, E., Gürtler, M., Pagliuca, F.W., et al. (2019). Charting cellular identity during human in vitro β-cell differentiation. Nature 569, 368–373. https://doi.org/10.1038/s41586-019-1168-5.

Wang, K., Wei, G., and Liu, D. (2012). CD19: a biomarker for B cell development, lymphoma diagnosis and therapy. Exp Hematol Oncol 1, 1–7. https://doi.org/10.1186/2162-3619-1-36.

Zhang, W., Jordan, K.R., Schulte, B., and Purev, E. (2018). Characterization of clinical grade CD19 chimeric antigen receptor T cells produced using automated CliniMACS Prodigy system. Drug Des Devel Ther 12, 3343–3356. https://doi.org/10.2147/DDDT.S175113.

Zhu, F., Shah, N., Xu, H., Schneider, D., Orentas, R., Dropulic, B., Hari, P., and Keever-Taylor, C.A. (2018). Closed-system manufacturing of CD19 and dual-targeted CD20/19 chimeric antigen receptor T cells using the CliniMACS Prodigy device at an academic medical center. Cytotherapy 20, 394–406. https://doi.org/10.1016/j.jcyt.2017.09.005.

